# Revealing the Ultrastructure of Live *Candida albicans* using Stimulated Emission Depletion Microscopy

**DOI:** 10.1101/2024.11.25.625149

**Authors:** Katherine J. Baxter, Liam M. Rooney, Shannan Foylan, Gwyn W. Gould, G. McConnell

## Abstract

*Candida albicans*, a commensal fungal pathogen, is a major cause of opportunistic infections in immunocompromised individuals. Understanding its cellular structures and pathogenic mechanisms is critical for developing targeted antifungal therapies. Stimulated emission depletion (STED) microscopy enables nanoscale visualization of cellular components, surpassing the diffraction limit of conventional light microscopy. In this study, we employed STED microscopy to investigate the ultrastructural organization of *C. albicans* in live specimens. We showed that dyes commonly used in STED microscopy of mammalian cells are ineffective for the study of *C. albicans*, and we showed the utility of Nile Red staining for visualising the organisation of dynamic cellular components, including tracking of lipid droplets, using time-lapse recording in experiments exceeding 12 hours. STED microscopy offered more than a two-fold improvement in resolution compared to confocal laser scanning microscopy applied to the same specimens with negligible photobleaching. This study demonstrates the utility of STED microscopy in advancing our understanding of *C. albicans* biology at the nanoscale, providing a platform for future investigations into fungal pathogenicity and antifungal development.

**GRAPHICAL ABSTRACT:** 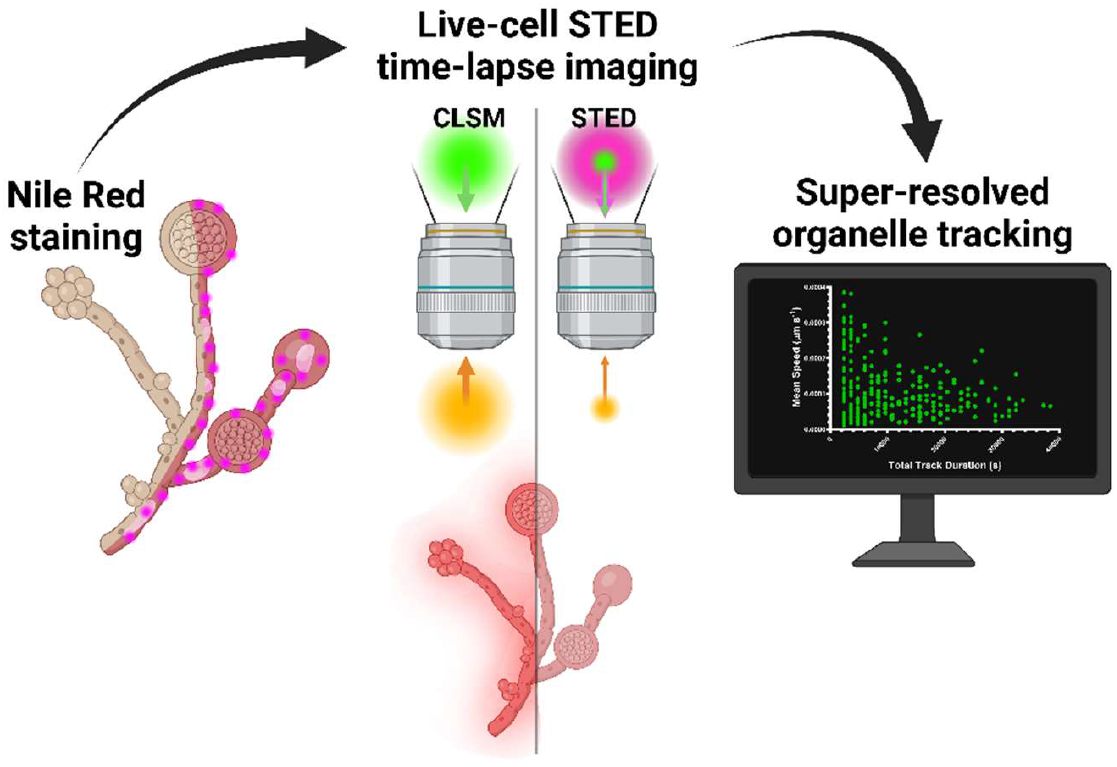

We present an optimised fluorescence staining method for super-resolution live-cell imaging of *Candida albicans*, using Stimulated Emission Depletion (STED) microscopy to resolve and track sub-cellular structures. We compare the performance of conventional confocal laser scanning microscopy (CLSM) to STED imaging, providing a three-fold resolution improvement beyond the diffraction limit. Finally, we perform live cell tracking to visualise and quantify the trajectories of multiple sub-diffraction limit-sized objects over a period of 12 hours, demonstrating the potential for live-cell STED imaging of *Candida* to visualise key processes involved in pathogenesis, drug resistance and infection.

## INTRODUCTION

*Candida albicans* is a pleomorphic fungus that plays a significant role in human health, particularly within microbial communities of the mouth, gut, and other mucosal surfaces^1^. As a part of the normal microbiota, *C. albicans* can exist harmlessly, aiding in nutrient processing and competing with harmful pathogens, which helps maintain a balanced microflora. However, *Candida* species are also opportunistic pathogens^1^; when the immune system is weakened or the host experiences dysbiosis (e.g., due to illness), *C. albicans* can lead to acute or invasive candidiasis^2^. These infections range from mild, superficial issues like oral thrush^3^ or fungal foot and nail infections^4^ to severe, systemic infections that can be life-threatening, particularly in immunocompromised individuals^5^.

One of the key critical structural features of *C. albicans* is its ability to switch between different forms, namely yeast, pseudohyphal, and hyphal, which allows it to spread and invade tissues effectively^6^. The complex cell wall of *C. albicans*, composed of β-glucans, chitin, and mannoproteins, provides protection and enables it to adhere to host cells, a necessary step for colonisation^7^. *C. albicans* can also form biofilms, dense layers of cells encased in a protective matrix, especially on medical devices, making these infections more persistent and resistant to antifungal treatments^8^. Additionally, the genetic adaptability and ability to secrete enzymes that break down host tissues enhance the virulence of *Candida* species^9^. These structural features make *C. albicans* a resilient pathogen, challenging to treat and control, especially in immunocompromised patients. Understanding these structural components is essential in developing effective antifungal therapies and preventive strategies.

Electron microscopy (EM) has proven indispensable for high-resolution imaging of *C. albicans* ultrastructure however it presents significant limitations, particularly in live-cell studies^10^. Standard EM techniques necessitate fixation, dehydration, and coating (for scanning EM^10^), or embedding and ultrathin sectioning (for transmission EM^11^), both of which inherently kill cells and preclude real-time analysis of critical dynamic phenomena, such as the study of lipid droplet trafficking. The chemical fixation and dehydration required can introduce preparation artefacts, potentially distorting cellular structures and misrepresenting the original architecture of the specimen. Furthermore, while EM provides exceptional structural resolution, images are only possible from a tiny area of the specimen, and images can lack molecular specificity, yielding only static, structural data without functional context. The requirement for a high-vacuum environment in most EM applications causes rapid desiccation and collapse of hydrated biological specimens, leading to structural aberrations that can also complicate accurate structural analysis. Moreover, the operational complexity and high costs also limit the feasibility of EM methods for large-scale studies, reducing the throughput. Although researchers can mitigate these limitations to some degree using complementary methods such as cryo-electron microscopy^12^ to preserve native cellular states via rapid cryofixation, these approaches also present technical challenges, underscoring the persistent obstacles in real-time ultrastructural analysis of *C. albicans*.

Fluorescence microscopy has significantly advanced our understanding of *C. albicans* by enabling the observation of cellular dynamics^13^, molecular interactions^14^, and pathogenic mechanisms^15^ in real-time and in a live-cell context. Unlike electron microscopy, fluorescence microscopy allows visualization of specific proteins, organelles, and other cellular components through fluorescent tagging, which has been instrumental in studying the morphological transitions between yeast, pseudohyphal, and hyphal forms of *C. albicans*^16^, providing critical insights into how cells adapt to environmental changes and invade host tissues. Furthermore, advances in fluorescent markers have enabled the study of intracellular signalling pathways, gene expression, and protein localization in *C. albicans*^17^.

Fluorescence microscopy has been invaluable for studying *Candida albicans*, but it does have notable limitations. Conventional fluorescence microscopy is limited by the diffraction of light, which restricts resolution to approximately 200–300 nm laterally and 500–700 nm axially. This makes it challenging to resolve the fine structural details of *C. albicans*, such as the formation of vacuoles, or tracking organelle movement.

Super-resolution microscopy methods are a family of powerful imaging technique that surpass the diffraction limit of light, enabling the observation of cellular structures at resolutions far beyond conventional light microscopy. Super-resolution methods can achieve resolutions as high as 20 nm, providing unprecedented detail of subcellular structures. Key super-resolution and single molecule localization microscopy techniques include stimulated emission depletion (STED)^18^, photoactivated localization microscopy (PALM)^19^, structured illumination microscopy (SIM)^20^, and stochastic optical reconstruction microscopy (STORM)^21^, each employing different approaches to break the diffraction barrier. STED microscopy was chosen here due to its high speed of imaging and compatibility with live cell imaging^22^. STED microscopy uses a pair of laser beams, one of which excites the fluorophores and another to deplete their fluorescence in a specific region, thus narrowing the effective excitation volume and achieving higher resolution^18^. The main benefit of STED microscopy is its ability to visualise nanoscale structures in live cells with high spatial and temporal resolution, making it particularly useful for studying dynamic processes such as protein localization, cell membrane organization, and cellular signalling. However, thus far, STED microscopy has not been widely used to study the ultrastructure of live *C. albicans*.

In this study, we have developed a robust protocol for live cell staining and time-lapse STED microscopy of *C. albicans*, enabling the visualization of dynamic cellular components, including organelles, specifically mitochondria and lipid droplet, distribution (Figure 1). The STED principle, shown in Figure 1(A), allowed us to resolve features beyond the diffraction limit of conventional light microscopy, offering new insights into the ultrastructure of *C. albicans* without the strenuous specimen preparation requirements of electron microscopy. Our approach highlights the potential of super-resolution imaging in fungal research, providing a clearer picture of the cellular machinery of *C. albicans* and advancing our understanding of the role this important pathogen plays in human health and disease.

**Figure 1.**
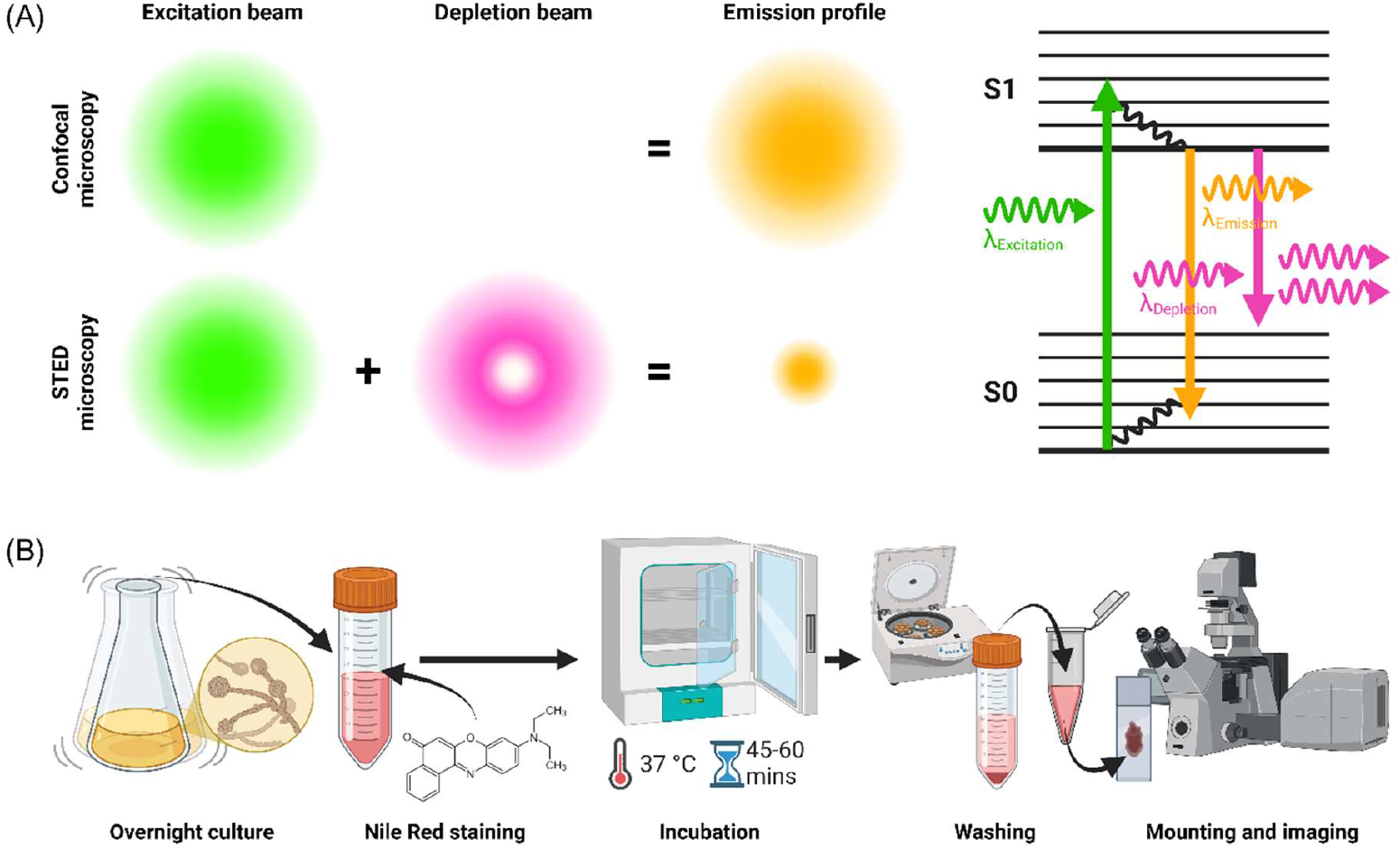
(A) A comparison of the emission profile acquired using confocal laser scanning microscopy (CLSM) and STED microscopy, along with a Perrin-Jablonski diagram describing the emission depletion during the fluorescence process. A depletion beam is aligned with the excitation laser during STED imaging to deplete the emission of fluorophores in the depletion region, resulting in a smaller overall emission profile and enhanced spatial resolution. Disruption of the internal conversion process causes relaxing electrons to immediately return from the excited state (S1) to the ground state (S0), releasing a red-shifted photon which is blocked by the microscope fluorescence emission filters, while fluorescence from S1 to S0 is collected from the confined excitation volume and used to form an image. (B) A methods workflow showing the growth, staining and preparation methods for live-cell STED imaging of *C. albicans*.

## METHODS

### Specimen preparation

To initiate *C. albicans* culture, a single colony was inoculated into 5 mL of Luria-Bertani (LB) medium supplemented with 0.2% glucose and incubated overnight at 37°C for approximately 16–18 hours. The duration of incubation influenced the extent of hyphal formation, with longer incubation times promoting more hyphal growth. After incubation, the culture was examined under a light microscope to assess the extent of hyphal development.

The effectiveness of multiple different stains for STED microscopy of live *C. albicans* were assessed, actin LIVE 460L (Abberior, Germany), DNA LIVE 590 (Abberior, Germany), and Nile Red (Sigma Aldrich, UK). These dyes were chosen because of their proven compatibility with STED microscopic imaging of mammalian cells^19,23,24^ using 488 nm and 561 nm excitation respectively and depletion at 775 nm. Both actin LIVE 460L and DNA LIVE 590 are optimised for live cell imaging. Nile Red is a commonly used lipophilic fluorescent stain, used for both live and fixed cell imaging^25^.

### Staining

Abberior LIVE stains were prepared as described on the manufacturer’s website (www.abberior.rocks), with lyophilised stains resuspended at a concentration of 1mM in 100% DMSO (Sigma Aldrich, UK). Immediately prior to use, LIVE stain stocks were diluted to a final concentration of 0.1 µM in 200 µL of fresh, pre-warmed LB supplemented with 0.2% glucose. To stain *C. albicans*, 200 µL of overnight culture in an Eppendorf tube was spun at 2500rpm for 2 minutes in a microfuge (Centrifuge 5418, Eppendorf) to pellet the cells, before resuspension in the 200 µL of pre-warmed LB containing 0.1 µM of stain. The Eppendorf tube containing cells and stain was then shielded from light and placed inside a universal tube for incubation at 37°C for 45 to 60 minutes at 220 rpm, prior to mounting.

For staining with Nile Red, 45 µL of PBS (pH 7.3) was added to a 55 µL aliquot of 90 µg/mL Nile Red in 100% Dimethylsulfoxide (DMSO) and mixed thoroughly, ensuring that the solution was kept in the dark to protect the dye from degradation. A 200 µL aliquot of the overnight *C. albicans* culture was transferred to an Eppendorf tube, followed by the addition of 20 µL of the Nile Red solution in 55% /45% DMSO/Phosphate Buffered Saline. The mixture was gently vortexed to ensure thorough mixing and to facilitate the staining of the cells. The Eppendorf tube was incubated as described above.

After incubation, Eppendorf tubes were removed from the universal tube and centrifuged (Centrifuge 5418, Eppendorf) at 2500 rpm for 2 minutes to pellet the cells. The supernatant, containing the staining solution, was carefully removed, and the pellet was resuspended gently in 200 µL of PBS (pH 7.3) to wash the cells and remove excess dye. An experimental workflow is presented in Figure 1(B). A minimum of three biological replicates were imaged for each live stain.

### Specimen mounting

For mounting the stained cells, a 4 µL volume of the resuspended cell sample was directly applied onto coverslips (22 mm × 22 mm, Thickness No. 1.5, VWR). Cells were immobilised by the addition of 8 µL of molten 0.5% agar spotted onto the cell sample, and immediately sandwiched with a glass slide to create a thin agar layer. Microscope slides were inverted onto the specimens for imaging.

### Microscopy

Imaging was performed using an inverted microscope equipped with confocal laser scanning microscopy (CLSM) and STED capability (STEDYCON, Abberior Instruments). Imaging was performed at room temperature, and a round-microscope enclosure prevented ambient light from reaching the specimen during imaging.

A 488 nm laser for excitation of specimens stained with LIVE 460L, and a 561 nm laser was used for excitation of fluorescence from both LIVE 590 and Nile red stained specimens while a third laser at a wavelength 775 nm performed the depletion needed for STED microscopy. Typical relative laser powers were approximately 2% at 561 nm for CLSM, and 10% for both the 488 nm and 561 nm lasers and 50% of the depletion laser for STED. These values were chosen to maximise imaging contrast at a relatively fast imaging speed. As all stains emitted fluorescence in the red region of the spectrum, a spectral detector spanning 575 nm to 625 nm was employed to minimise the contribution of backscattered laser light and improve image contrast for imaging of both LIVE stains and Nile red stained specimens.

Images were acquired using a high numerical aperture (NA) oil immersion objective (100x, NA 1.4), with a 60 µm diameter pinhole. The theoretical lateral resolution (r) of a CLSM^26^ is calculated from r=0.61λ/NA to be 268 nm for an emission wavelength (λ) of 616 nm and an objective lens with NA = 1.4.

Images were obtained with a pixel size of 30 nm to satisfy the Shannon-Nyquist sampling criterion^27^, with an image size of 100 µm × 80 µm, and a 10 µs pixel dwell time, with each image taking approximately 90 seconds to acquire. For time-lapse imaging, a 20-minute interval between images was chosen, and images were recorded over a 12-hour period. Images were acquired using STED microscopy and CLSM alternatively. Images were saved in 16-bit format using the proprietary .obf file type of the microscope.

### Image processing

Images in .obf format were processed using FIJI (v1.54f)^28^. In some time-lapse recordings, the specimens drifted in the lateral plane during imaging: for these data a manual drift correction was performed using FIJI with the “*Correct 3D drift*” in-built plugin^29^. Default settings were applied except the number of pixels of x and y correction were set to a maximum shift of 30 pixels each, and data were corrected in in the x and y direction only. The contrast of CLSM datasets was adjusted for the purpose of presentation using CLAHE^30^, with default settings applied. No contrast adjustment was performed to STED microscopy datasets. Image data were converted to OME TIFF format for study, and time-lapse recordings were resized and saved as .AVI files for display of movies.

### Image data analysis

To measure the resolution of our images we adopted two different methods. The first was to take a line ROI across the same region of a CLSM micrograph and a STED image, and to measure the separation between two finely spaced objects. However, this method introduces human bias. As such, we also applied Image Decorrelation Analysis (IDA), which is based on partial phase correlation and does not rely on user-defined parameters, and assesses the resolution from the whole image^31^. The FIJI plugin “*Image Decorrelation Analysis*” was used with default settings, using input data from both CLSM and STED microscopy. IDA returns the spatial frequency limit, which corresponds to the image resolution.

Lipid droplet trafficking was tracked using TrackMate^32,33^, an openly available plugin for FIJI. Lipid droplets were detected using a *Laplacian of Gaussian* filter with an estimated object diameter limit of 0.35 µm, a quality threshold of 1966, and sub-pixel localisation enabled, which proved effective in selecting liposome foci. A *Linear Assignment Problem* framework^34^ was used to track the foci throughout the time series. The maximum distance for frame-to-frame linking was set to 0.6 µm. The maximum gap closing distance was set to 0.6 µm, with a maximum frame gap of 2 frames. The resulting tracks were filtered to include tracks with a linearity of forward progression score below 0.9 for further analysis, removing perfectly linear tracks that likely arose due to errant combined tracks from closely neighbouring tracks. Tracks were presented with a colour coding corresponding to the track duration and the track features were exported as .CSV files and transcribed for plotting and statistical analyses using GraphPad Prism (v8.0.2; Mathworks). Tracking and analysis were performed using a 64-bit Windows Server 2016 standard operating system (v.1607) with two Intel® Xeon® Silver 4114 CPU processors at 2.20 and 2.19 GHz and 1.0 TB installed RAM.

## RESULTS

### Dyes suitable for STED microscopy of mammalian cells are ineffective for labelling of *C. albicans*

Figure 2 shows images of *C. albicans* labelled with Nile Red obtained with (A) CLSM and (D) STED microscopy. Figures 2(B) and 2(E) show CLSM and STED data obtained from *C. albicans* labelled with actin LIVE 460L, and Figures 2(C) and 2(F) also show CLSM and STED data respectively, here with *C. albicans* labelled with DNA LIVE 590.

**Figure 2.**
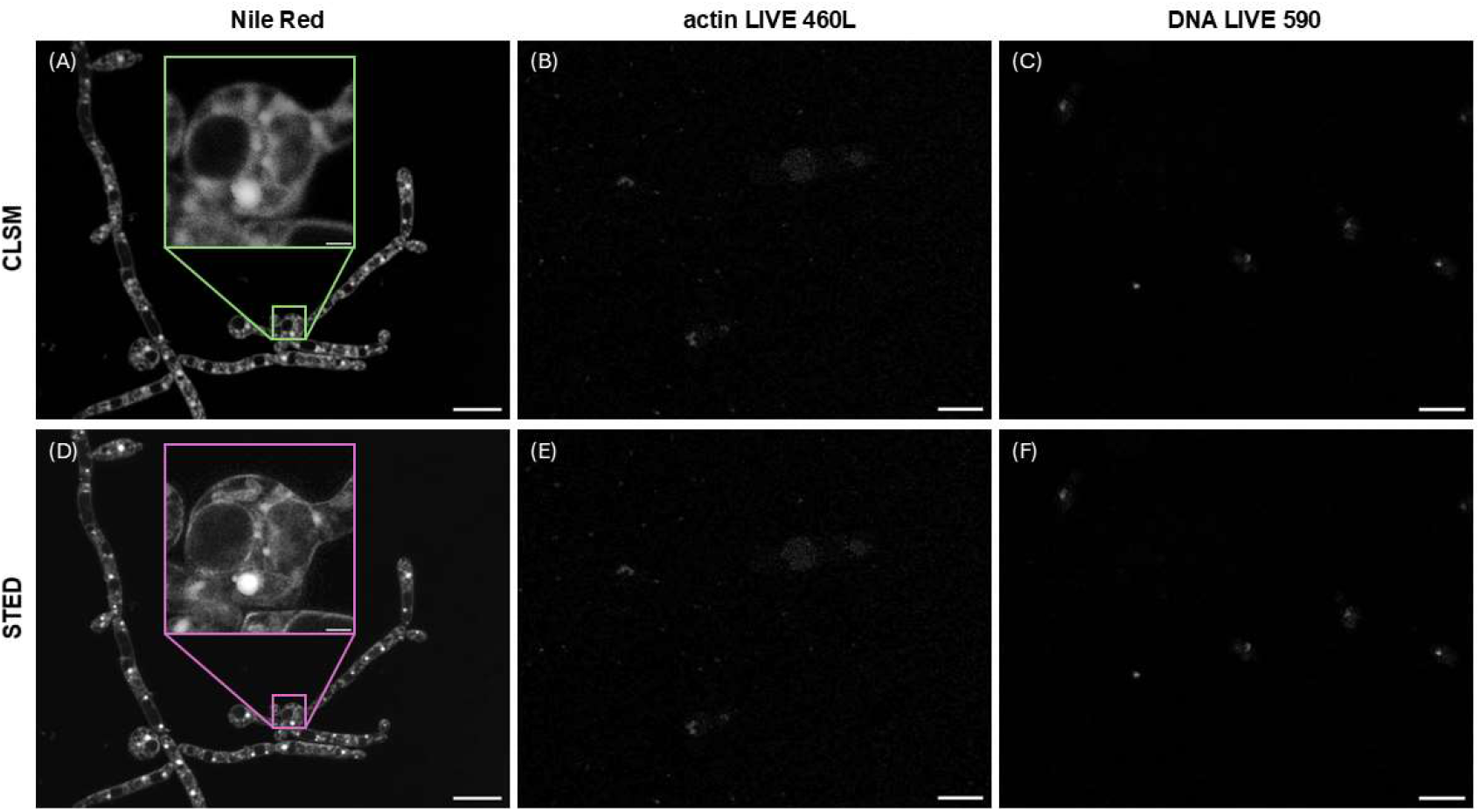
(A) *C. albicans* labelled with Nile Red, imaged with a CLSM. A digitally magnified ROI is shown with a green box, confirming intracellular staining. (B) *C. albicans* labelled with actin LIVE 460L, imaged with a CLSM. (C) *C. albicans* labelled with DNA LIVE 590, imaged with a CLSM. (D), (E), and (F) show the STED counterparts of (A), (B), and (C). A digitally magnified ROI is shown in (D), showing the level of detail improvement possible with STED microscopy compared to CLSM.

Fluorescence from the specimens labelled with Nile Red is clearly visible. Digitally magnified regions of interest (ROIs) are shown in the CLSM data in Figure 2(A) and in the STED image data presented in 2(D): in both cases, it is evident that Nile Red stained not only the cell boundary but also intracellular structures, including brightly stained structures representing lipid droplets. By comparison, virtually no fluorescence signal was visible in specimens labelled with either actin LIVE 460L or DNA LIVE 590.

### STED microscopy reveals ultrastructure of live *C. albicans* in specimens stained with Nile Red

Figure 3 shows representative images of *C. albicans* stained with Nile Red imaged with CLSM and STED microscopy. Figure 3(A) shows a CLSM image, and Figure 3(B) shows the equivalent STED microscope image. Two ROIs are highlighted with cyan and yellow boxes, and digital zooms of the CLSM and STED microscope images are shown in (C) & (E) and in (D) & (F), respectively. There is a slight temporal offset of around 1 minute between the CLSM and STED microscope images because they are acquired sequentially, which manifests as a slight disparity in the image content between the CLSM and STED micrographs, but with most of the structures retained.

**Figure 3.**
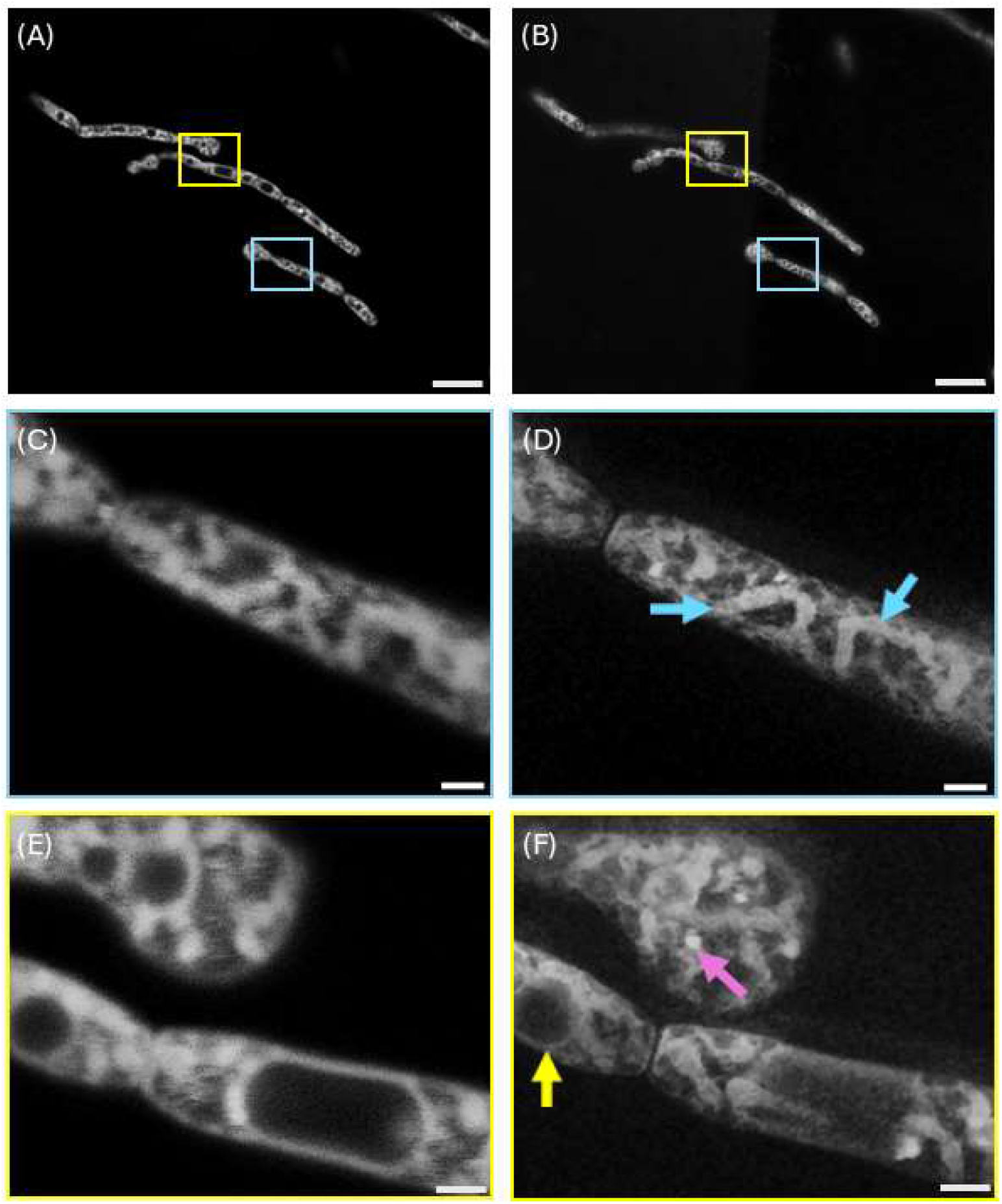
CLSM and STED microscopy of live *C. albicans* stained with Nile Red. (A) Representative confocal microscopy image. (B) Representative STED microscopy image of the same field of view as shown in (A). Cyan and yellow boxes highlight regions of interest (ROIs). Digitally magnified ROIs of CLSM data are shown in (C) and (E), while digitally magnified ROIs of STED data are shown in (D) and (F). Cyan arrows in (D) show mitochondria with the cristae visible, yellow arrow in (F) shows a sub-micron diameter vacuole interacting with mitochondria, and the magenta arrow in (F) highlights an example of a lipid-rich vesicle. Features indicated by arrows are not clearly visible in the corresponding CLSM micrographs (C) and (E). Scale bars for (A) and (B) = 10 µm, scale bars for (C) to (E) = 1 µm.

A clear resolution improvement is shown in the STED microscopy images. Figure 2(D) shows the lipid-rich elongated mitochondria^35^ clearly in the STED data, with the cristae visible: the mitochondria are difficult to discriminate from other cellular structures in the corresponding CLSM data presented in Figure 2(C).

The STED data presented in Figure 3(F) also reveal ultrastructure that is not visible in the confocal microscopy counterpart, Figure 3(E). Liposomes are visible: an example is highlighted with a magenta arrow in Figure 3(F). Membrane-bound vacuoles interacting with mitochondria are also observed, as indicated by the yellow arrow in Figure 3(F). These sub-diffraction limit-sized structures are not clearly visible in the CLSM data.

### STED microscopy of *C. albicans* offers more than a two-fold resolution improvement compared to CLSM

Resolution measurements of CLSM and STED data confirm the improvement in resolution using the STED imaging approach. CLSM and STED images of *C. albicans* labelled with Nile Red are shown in Figure 4(A) and 4(D) respectively, and a line ROI is shown in 4(A). The same ROI was applied to 4(D) for the purpose of measuring the resolution of the image. As shown in Figure 4(B) it was not possible to resolve the structures highlighted in 4(A). However, it was possible to resolve the discrete membranous structures in 4(D), and the simple line ROI measurement produced a resolution measurement of 120 nm from the STED data, as shown in 4(E). A direct comparison of the resolution of CLSM and STED microscopy is possible using IDA. The resolution of the CLSM and STED images shown in Figures 4(A) and 4(B) was measured to be 417 nm and 136 nm respectively, using IDA. While the resolution of the CLSM was measured to be poorer than predicted by the theoretical limit of diffraction, the STED microscope offered approximately a factor of two improvement in resolution compared to the theoretically best possible resolution of the CLSM^26^, and more than a three-fold improvement when comparing practical CLSM and STED microscopy, as shown in Figures 4(C) and 4(F).

**Figure 4.**
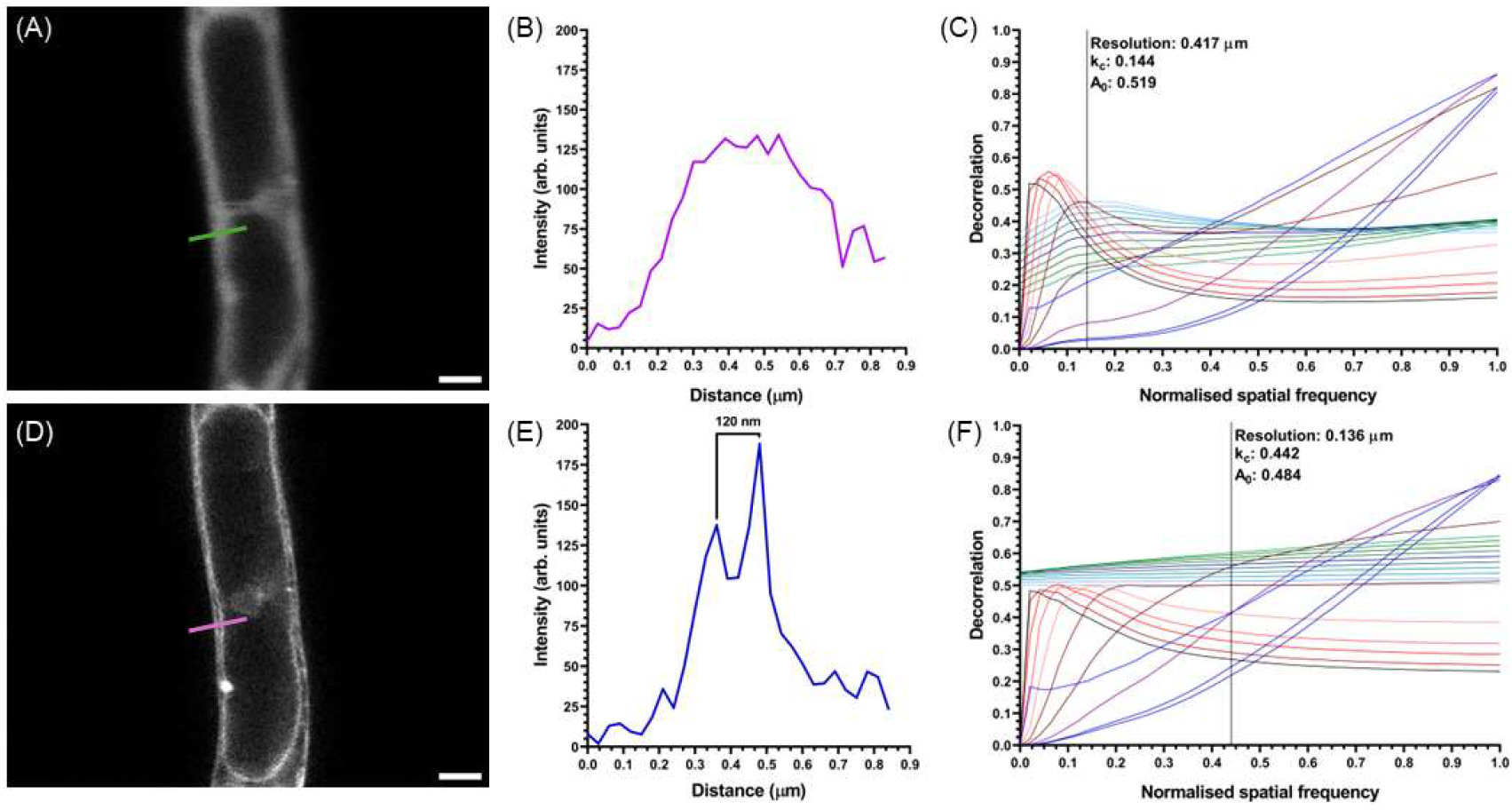
(A) CLSM image of *C. albicans* labelled with Nile Red. A green line shows a ROI. (B) Line intensity profile of the ROI shown in (A). (C) Image decorrelation analysis of (A) gives a resolution of 417 nm for this CLSM micrograph. (D) STED microscopy image of *C. albicans* labelled with Nile Red, showing the same ROI as (A) with a magenta line. (E) Line intensity profile of the ROI shown in (D) for the STED data. (F) Image decorrelation analysis of (D) gives a resolution of 136 nm for these STED data. Scale bars for (A) and (D) = 1 µm.

A representative time-lapse recording of Nile Red stained *C. albicans* is presented as Movie 1. Both the CLSM and STED microscopy datasets showed organelle motility throughout the full duration of the 12-hour recording, indicating sustained cell viability. However, only the high-resolution detail afforded by STED microscopy offered the possibility to measure and track of sub-diffraction limited intracellular structures. Minimal photobleaching was observed in these long-term recordings. A small amount of specimen drift in the axial direction was observed from t=540 minutes in one part of imaged field, leading to a decrease in fluorescence signal and blurring of this region of the image, but the other cells in the specimen remained in focus throughout the full duration of the recording.

Figure 5 shows the tracking and quantification of lipid droplet trafficking in live *C. albicans* over 12 hours acquired using STED microscopy. Tracks are presented in Figure 5(A), colour-coded by track duration. Lipid droplet trafficking was observed in multiple cells throughout the acquisition, with a mean track duration of 180 minutes (Standard Error in the Mean (SEM) = 8 minutes 47 seconds, Standard Deviation (SD) = 141 minutes 14 seconds). Figure 5(B) shows the relationship between lipid droplet trafficking speed and total distance travelled. In general, the mean speed of a given liposome trafficking event scaled with the length of the track, meaning longer tracks were typically slower than short tracks. The mean speed of lipid droplet trafficking was measured at 110.0 nm s^-1^ (SEM = 5.2 nm s^-1^, SD = 83.0 nm s^-1^), with a maximum total distance of 5.82 µm. Movie 2 shows the lipid droplet trafficking dynamics in a digitally magnified ROI from Figure 5, overlayed with tracks colour coded as described for Figure 5. The sub-cellular ultrastructure and organelle motility can be observed over an excerpt of 650 minutes from the full 12-hour time series. In addition to the tracked lipid droplet dynamics, vacuole migration, and mitochondrial motility can also be clearly resolved by live-cell STED imaging but could not be seen in the CLSM data.

**Figure 5.**
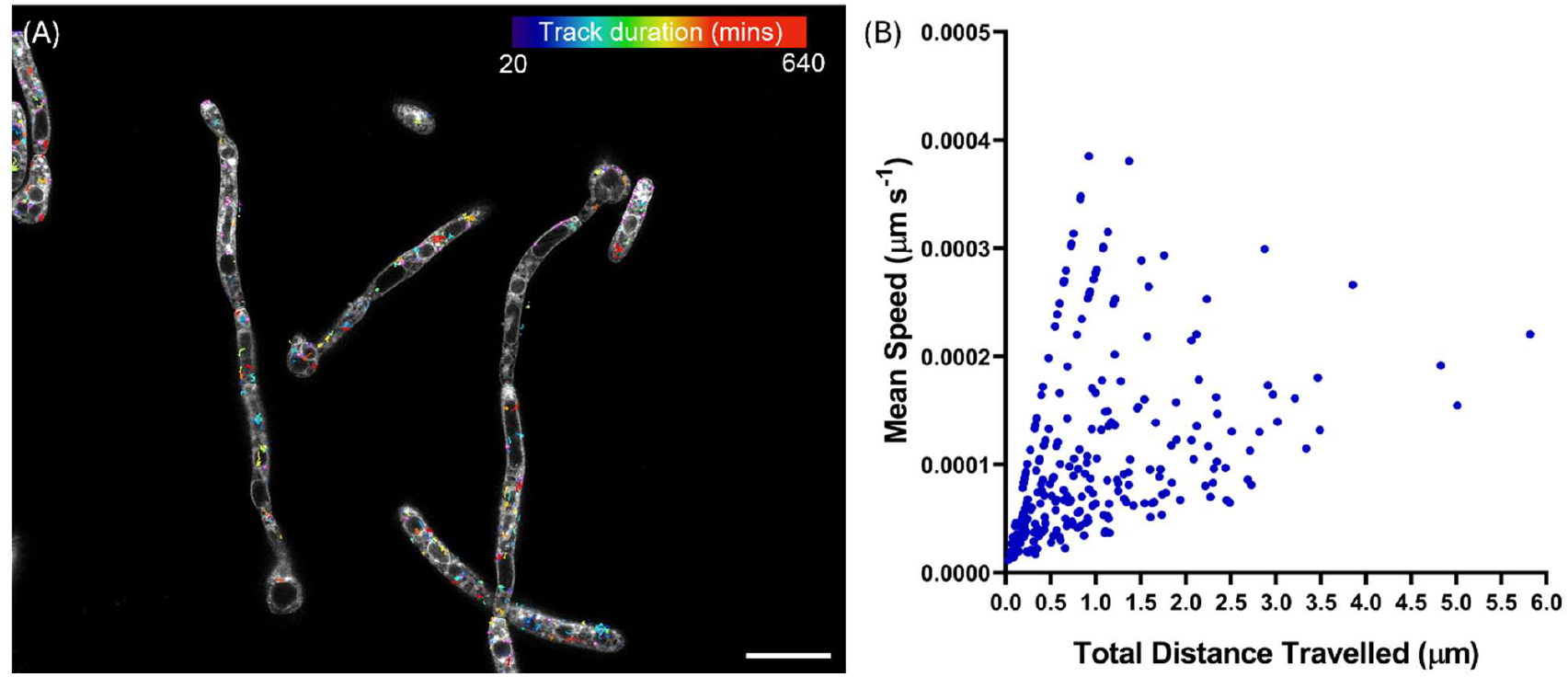
(A) An overlay of a STED image of *C. albicans* labelled with Nile Red and the tracks of lipid droplet trafficking over 12 hours, with tracks colour-coded according to the total track duration. (B) A plot showing the mean speed versus the total distance travelled by each liposome during the time series shown in (A). Scale bar = 10 µm.

## DISCUSSION

In this study, we employed STED microscopy to investigate the ultrastructure of live *C. albicans*. Our findings provide super-resolution insights into the dynamic organisation of the mitochondria and their interaction with other organelles, including lipid droplets. The dynamic and regulated association between mitochondria and lipid droplets in mammalian cells is tightly linked to metabolic status^36^, and thus our observation of close apposition of lipid droplets and mitochondria in *C. albicans* suggest that this is an area worthy of further exploration.

We observed that some fluorescent labels designed for STED microscopy, while effective for the study of mammalian cells, were not suitable for this super-resolution optical imaging method when applied to *C. albicans*. Both LIVE stains, like Nile Red, have no charge^37^, so their ineffectiveness in *C. albicans* is unlikely to be caused by polarity. The molecular weight of LIVE 460L and LIVE 590 are not published, but based on the STAR derivative structure, which is approximately 880 Daltons^38^ they are likely to be larger than that of Nile Red, which is a fluorescent molecule with one of the lowest molecular weights (318 Daltons)^39^. The cell wall of *C. albicans*, which is around an order of magnitude thicker than the plasma membrane of mammalian cells^7^, may be preventing the ingress of LIVE 460L and LIVE 590.

Our work with Nile Red has proven the benefit of STED microscopy for studying the living ultrastructure of *C. albicans*, but this staining method labels only lipids, including lipid bilayers and, notably in the context of this study, intracellular lipid droplets. Commonly used STED dyes at similar excitation wavelengths to those used in our study such as ATTO 647N, DyLight 650, and Alexa Fluor 647 have comparatively high molecular weights at 660 Daltons, 620 Daltons, and 1100 Daltons respectively. To take advantage of the molecularly selective affinity that fluorescent labelling confers, e.g. for multi-colour STED microscopy of organelles labelled with protein-specific dyes, alternative fluorescent dyes with lower molecular weights compatible with STED imaging may be needed. Such new stains could open the possibility of studying other dynamic cellular processes in *C. albicans*. These include the study of cell division during morphogenesis, enabling the observation of septin filaments at the site of cell division^40^, understanding the remodelling of vacuoles in response to environmental stressors or changes in nutrient availability^41^, the role of extracellular vesicles in the modulation of host-pathogen interactions^42^, and unveiling how adhesin expression changes in *C. albicans* during the formation of biofilms on the surface of medical devices^43^. There is also considerable potential to apply this methodology to the study of drug and toxin effector molecules, with improved prospects for characterising how antifungal agents disrupt cellular processes^44^.

While the 120 nm resolution of this imaging method offers a considerable improvement over conventional technologies such as CLSM, a much higher spatial resolution is theoretically possible with STED microscopy. The theoretical resolution of STED microscopy is primarily determined by the wavelength of the excitation light, the size of the fluorescence emitting region, and the intensity and size of the depletion beam, and can provide resolutions of between 20 nm to 50 nm^45^. However, to reach these very high resolutions, specific experimental conditions must be met. These include using a very high depletion laser power, which can be damaging to the specimen and can prohibit live cell imaging. For single-shot imaging, it may be possible to improve our reported resolution considerably using higher optical powers. It is possible that the relatively thick cell wall in *C. albicans*^7^ is also scattering the ordinarily well-defined laser spots, causing defocus and a decrease in measured resolution. A possible solution – which has already been proven in the use of STED microscopy to the study of mammalian cell specimens – is the use of adaptive optics (AO). AO systems typically use deformable optical elements to dynamically adjust the optical path and correct distortions^46^. This correction helps maintain the high resolution achieved by STED microscopy^47^, even in thicker or more scattering samples (e.g., tissues or complex biological environments), where traditional methods may struggle. This, however, increases both the cost and complexity of the instrumentation, and AO elements may require on-going calibration and optimisation, which may not be compatible with live cell studies. SIM^20^ may offer an alternative method to STED for super-resolution imaging of *C. albicans*, but in this method a minimum of nine images are required to improve the resolution by a factor of two, and this method is relatively slow to capture data and can cause photobleaching.

STED microscopy enabled tracking of intracellular organelles with high precision and minimal photobleaching over more than 12 hours. Our data resolved organelle movement, in this case lipid droplets, occurring over distances of less than 2 µm or within tight confinement neighbourhoods. Such short trajectories would be almost impossible to measure with an CLSM because of the relatively poor resolution afforded by diffraction limited optics. The measured track distances broadly agree with the scale of *C. albicans* cells and previously reported velocimetry studies in mammalian cell biology^48,49^. Lipid droplet, vesicle, and vacuole trafficking have been reported to play multiple roles in the virulence, pathogenesis, drug resistance, and ageing of *Candida* species. Vacuolar dynamics are highly regulated during normal development and physiology, maintaining ion homeostasis and promoting hyphal formation^50^, with vacuole-mediated autophagy being linked with pathogenesis^51^. Moreover, lipid droplet formation has been well documented in yeasts, showing their accumulation over the lifespan of the organism^52^,^53^ and vertical transfer through lineages of daughter cells^54^. However, the ability to track lipid droplets and vacuole dynamics over long periods has been hampered due to the inability to resolve such small structures in conventional diffraction-limited imaging. By providing means to resolve these organelles in both space and time, researchers may adapt our methods to study the role of lipids in pathogenesis and drug resistance; for example, in understanding dynamic morphological changes in the organisation of the *Candida* vacuole that promote yeast-to-hyphae transitions^51^, or in maintaining the specific protein and lipid composition of the membrane, facilitating tissue invasion and infection. Lipid droplet dynamics are also inherently linked to drug resistance and antimicrobial resistance. The punctate lipid droplet distribution that we observed has been reported to disperse because of flippase transporter mutations, such as in Dsr2^55^ which is essential for filamentous growth, phosphatidylserine and ergosterol distribution, and maintaining copper and fluconazole resistance. Other studies have extended to the documentation of numerous mechanisms by which lipids promote drug resistance^56^, for example by trapping and occluding toxins in lipid droplets^57^, thus minimising the efficacy of antifungal treatments.

Our results show the benefits of 2D STED for super-resolved imaging of *C. albicans*, but 3D STED is also possible^58^. 3D STED microscopy offers significant benefits over 2D STED microscopy, particularly for studying complex three-dimensional structures within biological samples. 3D STED would offer insights into hyphal formation by providing high-resolution imaging of the hyphal tip^59^, visualisation of protein trafficking^60^, and the distribution of signalling molecules during cell adhesion^61^. However, 3D STED has considerable limitations, including high rates of photobleaching and phototoxicity that can compromise live cell studies. Moreover, the additional time required to scan in the axial direction for volumetric imaging may lead to difficulties in object tracking.

While we have demonstrated our method in *C. albicans*, our work raises the possibility to apply STED microscopy to other *Candida* species, such as *Candida glabrata* or *C. auris. C. glabrata* exhibits resistance to common anti-fungal treatments (e.g. fluconazole)^63^, and *C. auris* is a healthcare-acquired multi-drug resistant pathogen recalcitrant to various anti-fungal treatments and decontamination methods^64^. Our method could be applied to investigate how drug resistance is mediated at the molecular level in both species, for instance, by examining changes in vacuole architecture or subcellular distribution of drug efflux pumps^65^. Moreover, our method could be adapted to the study of polymicrobial specimens and interkingdom interactions, such as those between *C. albicans* and bacteria including *Staphylococcus aureus*, e.g. to identify localised regions of signalling molecule accumulation, e.g. farnesol, within co-cultures^66^. This would provide valuable insight into how *C. albicans* communicates with other microbes or how it modulates its own behaviour and virulence in response to signals from other species.

## MOVIE CAPTIONS

**Movie 1**. Representative time-lapse confocal laser scanning microscopy and stimulated emission depletion microscopy (STED) imaging of *Candida albicans* stained with Nile Red.

**Movie 2**. Representative time-lapse stimulated emission depletion microscopy (STED) imaging of *Candida albicans* stained with Nile Red showing trafficking of lipid droplets over 12 hours. Tracks are colour coded according to the total track duration.

## AUTHOR CONTRIBUTIONS

Conceptualization: KJB, GM; Methodology: KJB, LMR, SF; Validation: KJB, LMR, SF, GM; Formal analysis: LMR, GM; Investigation: KJB, GM; Resources: GM, GWG; Data curation: KJB, LMR, GM; Writing - original draft: GM, LMR; Writing - review & editing: KJB, LMR, SF, GWG, GM; Visualization: LMR, GM; Supervision: GM, GWG; Project administration: GM; Funding acquisition: GM, GWG.

## FUNDING

KJB was supported by an EPSRC Impact Acceleration Account awarded to the University of Strathclyde (EP/X525820/1). LMR was supported by the Leverhulme Trust. GWG and GM were supported in part by the Biotechnology and Biological Sciences Research Council (BB/V019643/1). SF, GWG, and GM were supported in part by the Biotechnology and Biological Sciences Research Council (BB/X005178/1). GM was supported by the Biotechnology and Biological Sciences Research Council (BB/T011602/1 and BB/W019032/1) and the Leverhulme Trust.

## CONFLICT OF INTEREST STATEMENT

The authors declare no competing interests.

## Notes

https://video.wixstatic.com/video/4bc574_d9e77b65590a4668acda2d73df257db6/1080p/mp4/file.mp4

https://video.wixstatic.com/video/4bc574_8afa219559a2457bb8db2fa66e4fa731/480p/mp4/file.mp4

### Competing Interest Statement

The authors have declared no competing interest.

